# Quantitative proteomic analysis of soil-grown *Brassica napus* responses to nutrient deficiency

**DOI:** 10.1101/2024.09.01.610712

**Authors:** LE Grubb, S Scandola, D Mehta, I Khodabocus, RG Uhrig

## Abstract

Macronutrients such as nitrogen (N), phosphorus (P), potassium (K), and sulphur (S) are critical for plant growth and development. Field-grown canola (*Brassica napus* L.) is supplemented with fertilizers to maximize plant productivity, while deficiency in these nutrients can cause significant yield loss. A holistic understanding of the interplay between these nutrient deficiency responses in a single study and canola cultivar is thus far lacking, hindering efforts to increase the nutrient use efficiency of this important oil seed crop. To address this, we performed a comparative quantitative proteomic analysis of both shoot and root tissue harvested from soil-grown canola plants experiencing either nitrogen, phosphorus, potassium, or sulphur deficiency. Our data provide critically needed insights into the shared and distinct molecular responses to macronutrient deficiencies in canola. Importantly, we find more conserved responses to the four different nutrient deficiencies in canola roots, with more distinct proteome changes in aboveground tissue. Our results establish a foundation for a more comprehensive understanding of the shared and distinct nutrient deficiency response mechanisms of canola plants and pave the way for future breeding efforts.

## 1 INTRODUCTION

Canola (*Brassica napus* L.) is one of the leading oil seed crops of high economic value for food, animal feed, and biofuel, second only to soybean as the most consumed vegetable oil globally^1^. As a result of conventional breeding efforts, canola oil consists of lower erucic acids and glucosinolates^2^, as well as high mono-unsaturated fatty acid and low poly-unsaturated fatty acid levels^3^, resulting in it being a healthier option for seed oil consumption. As a result, there is increasing demand for canola oil products, driving research and breeding efforts for improved seed yield and oil production^4,5^. The overall success of these efforts has resulted in a >450-fold increase in seed yield since 1978^6^. However, the emergence of biotic and abiotic production challenges, especially due to climate change, have further increased the need to develop canola varieties resistant to disease and abiotic stresses. In particular, the biotic stresses of sclerotinia stem rot (*Sclerotinia sclerotiorum*)^7^, clubroot (*Plasmodiophora brassicae*)^8^, and blackleg (*Leptosphaeria maculans*)^9,10^, as well as abiotic stress such as heat^11,12^ and drought^13^ are some of the major threats to canola production on a global scale. Beyond these stresses, an often-overlooked component of the canola production landscape is the need for large-scale fertilizer application to sustain high-yielding crops from year-to-year. Despite the positive impacts on crop yield, excessive fertilizer application can have wide-ranging environmental impacts, while also imposing significant costs on producers. On average, approximately half of the fertilizer that is applied is effectively taken up and used by plants, leading to agricultural runoff resulting in eutrophication within aquatic ecosystems^14,15^, and the production of the greenhouse gas nitrous oxide by soil microbes^16^. Given the combination of protracted timelines for new trait introgression into elite, commercially grown germplasm, and the increasing costs and decreasing availability of high intensity fertilizers into the future, understanding how canola responds to nutrient stress at the molecular level now will contribute to future sustainability in canola production.

Nitrogen (N), phosphorus (P), potassium (K) and sulphur (S) are four of the essential macronutrients required for plant growth and development^17^. Deficiencies in any of these nutrients can have dramatic impacts on plant growth and yield. Soil N is taken up by plants in the form of nitrate (NO_3_^-^) or ammonium (NH_4_^+^). N plays a key role in plant metabolism, as a key constituent of many organic compounds such as proteins and nucleic acids, as well as chlorophyll, co-enzymes, phytohormones and secondary metabolites^18^. P is indispensable for plant growth and metabolism, acting as the main component of many biomolecules such as DNA, RNA, ATP, NADPH, and phospholipids, as well as playing a major role in photosynthesis and respiration^19^. P uptake from the soil occurs in the form of inorganic phosphate (orthophosphate; Pi), which can be limited by accessibility as Pi tends to form complexes with metal ions^20,21^. K is taken up in its ionic K^+^ form, in which it remains following uptake and functions as a major osmolyte and positive charge source for counterbalance, maintaining electrical homeostasis and enzyme activation^22^. S is taken up in the form of sulfate and is incorporated into the amino acids cysteine (Cys) and methionine (Met), which are major constituents of many enzymes, structural proteins as well as metabolic proteins^23^. S is also a component of glucosinolates, which act as key players in defense signaling, and serves as a base component of the plant hormones ethylene and abscisic acid (ABA), which are important for plant growth and development^24^. In comparison to wheat (*Triticum aestivum*), canola has a high nitrogen uptake but low recovery in reproductive tissues, with much lost to the soil in dead leaves during the growth cycle, therefore requiring proportionally more nitrogen input^25^. Additionally, with critical components of high-energy fertilizers, such as phosphate, becoming limiting in the next 100 years^26^, there is a need to develop canola varieties with higher nutrient use efficiency (NUE). Correspondingly, increasing NUE requires a robust systematic understanding of how plants adapt to nutrient deprivation in order to identify targets for breeding and/or biotechnology efforts.

To date, there has been limited systematic examination of multiple nutrient deficiencies in canola in a single experiment, with most studies focusing on identifying starvation responses at either the phenotypic and/or transcriptomic level, or by performing genome-scale informatics analyses of nutrient uptake and signaling gene families^27–36^. Such systems-level analyses include: studies examining ammonium/nitrate^37–44^, phosphate^44–48^, potassium^44,49–53^, and sulphur^54–57^ deficiency. However, no studies to date have examined deficiencies in multiple nutrients in a single study using the same canola variety at the proteomic level. As it is well established that transcriptome analyses only approximate cellular state and function across organisms^58–63^, it is essential that we start to define how canola responds at the protein level. Importantly, the majority of previous studies have focused on whole-plant, rather than organ-level analyses, and many were performed using hydroponic conditions, which is distinct from soil-grown experimentation.

Much of what we know about canola biology at the systems-level is based on transcriptomic technologies^64,65^, or conventional low-resolution two-dimensional gel electrophoresis experiments^65,66^. However, over the past several years, more advanced mass spectrometry technologies and quantitative proteomic acquisition workflows have begun to emerge that now allow for more robust deep, quantitative proteome profiling. Typically, these quantitative proteomics workflows have combined data dependent acquisition (DDA) with isobaric label tagging (e.g. TMT) and extensive offline fractionation to obtain deep protein profiling. This has included studies examining seed development^67^, response to waterlogging^68^, chloroplasts^69^, leaf rolling^70^, heavy metal toxicity^71^, the trade-off between seed protein and oil content^72^, and adaptive responses to nutrient deficiencies^73,74^, amongst others. However, sample fractionation followed by DDA acquisition results in increased instances of missing values and bias towards highly-abundant proteins relative to other quantitative approaches^75^, which can limit biological insights and decrease the generation of actionable data for new breeding opportunities. Here however, we apply a new label-free data independent acquisition (DIA) approach called BoxCarDIA^76,77^ to quantify changes in soil-grown canola (var. Westar) shoot and root proteomes in response to multiple nutrient stress conditions: -N, -P, -K, and -S.

Owing to its high seed yield, high oil content and early maturity^78,79^, we utilized the license-free Westar canola cultivar as our canola model, as it is ideal for molecular characterization and experimentation. Collectively, the results generated by this study provide a robust resource for understanding nutrient responses in canola, in addition to providing critically needed insight into the key proteins and cellular processes modulated in canola shoots and roots in response to nutrient deficiencies, offering actionable data for immediate implementation.

## 2. MATERIALS AND METHODS

### 2.1 Plant growth and tissue harvesting

Canola seeds (*Brassica napus* cv. Westar) were sterilized for 2 min in 70% ethanol, followed by a 7 min treatment in 30% bleach solution (Chlorox® 7.5%) and subsequently rinsed three times with sterile water. Seeds were then placed on media plates containing ½ Murashige and Skoog (MS), 1% sucrose and 7 g/L agar (Phytotech) at pH 5.8. Following 3 days stratification at 4 °C in the dark, the seeds were put to germinate under a 12 h light at 21 °C and 12 h dark at 19 °C photoperiod. Germinated seedlings were transferred to a soil mixture ProMix (PRO-MIX® BY MYCORRHIZAE™): Perlite (2:1) (adapted from Vishwanath et al. (2013)^80^ and grown under the same light and temperature regime as described above.

No fertilizer was added at transplantation or during the next 2 weeks of cultivation. When necessary, 1 L Milli-Q water was added per tray containing 4 canola plants in individual pots. After two weeks of starvation, treatment started by adding 2 L per tray of Hoagland solution complete or deplete of nitrogen (-N), phosphate (-P), potassium (-K) or sulphur (-S) according to ^81^ (see **SuppTable 1**). After 11 days of treatment, shoot and root tissue was harvested and directly flash frozen in liquid nitrogen prior to extraction.

**Table 1.** Quantified and significantly changing proteome (Log2FC>0.58, q<0.05) for roots and shoots under -N, -P, -K and -S.

|  | Quantified Proteins |  |  |  |  | Significantly Changing Proteins<br>(Log2FC>0.58; q-value ≤ 0.05) |  |  |  |  |
| --- | --- | --- | --- | --- | --- | --- | --- | --- | --- | --- |
|  | Control | N | P | K | S | Total | N | P | K | S |
| Shoots | 4386 | 4378 | 4448 | 4468 | 4365 | 2200 | 1471 | 568 | 755 | 508 |
| Roots | 1169 | 1771 | 1592 | 1486 | 1493 | 713 | 350 | 371 | 357 | 298 |

### 2.2 Elemental tissue analysis

Approximately 3 g of freshly frozen canola shoot tissue was ground to a fine powder and freeze-dried for 24 h. Samples were sent to and processed by the Natural Resources Analytical Laboratory (NRAL) of the University of Alberta (Canada). A Thermo iCAP6300 Duo ICP-OES instrument was used to analyze total Na, K, Ca, Mg, Fe, B, Mn, Zn, Cu, Mo, Ni, S and P. Samples were digested with HNO_3_ and atomized at high temperature (5500 to 8000 K). Emission patterns unique to each compound were then detected by the optical emission spectrometer. Multi-element and separate certified standard solutions (SCP Scientific) were used for calibration and as external reference standards.

Total N and C was analyzed with a ThermoScientific, Flash 2000 Organic Elemental Analyzer. Samples were weighed in open silver capsules and acidified with sequential additions of 50 µL of 1 M HCl to remove inorganic C. HCl was added until no further reaction was observed, as indicated by bubbles formed by the reaction of HCl and carbonate. The samples were then oven dried at 70 °C overnight and sealed. Samples were then analyzed by combustion elemental analysis for % C as described below. Complete combustion of the sample was achieved by dropping a known mass of sample, packed into a tin or silver capsule, into a combustion tube containing chromium (III) oxide and silvered cobaltous/cobaltic oxide catalysts. An aliquot of purified oxygen was added to the quartz tube generating a flash combustion reaction, increasing the reaction temperature from 1020 °C to 1800-2000 °C. The C in the sample was converted to CO_2_, and the N was converted to N_2_ and NO_x_. These combustion gases were carried via UHP helium through a reduction furnace filled with reduced copper wires, reducing NO_x_ species to N_2_. The gas stream passed through two sorbent traps to remove water, magnesium perchlorate for N_2_ and EMASorb, (formerly Carbasorb) for CO_2_ analysis. The resulting N_2_ gas and/or CO_2_ gas was separated on a 2 m x 6 mm OD stainless steel Porapak QS 80/100 mesh packed chromatographic column and detected quantitatively by a Thermal Conductivity Detector (TCD). The integrated TCD peak signal in the resulting chromatogram is directly proportional to the amount of C and N present in the sample which, along with the sample weight, is used to calculate %C and %N (w/w).

### 2.3 Tissue processing, protein extraction and digestion

Quick-frozen canola shoots and roots were ground to a fine powder under liquid N_2_ using a mortar and pestle. Ground samples were aliquoted into 400 mg fractions. Aliquoted samples were then extracted at a 1:2 (w/v) ratio with a solution of 50 mM HEPES-KOH pH 8.0, 50 mM NaCl, and 4% (w/v) SDS. Samples were then vortexed and placed in a 95 °C table-top shaking incubator (Eppendorf) at 1100 RPM for 15 min, followed by an additional 15 min shaking at room temperature. All samples were then spun at 20,000 x g for 5 min to clarify extractions, with the supernatant retained in fresh 1.5 mL Eppendorf tubes. Sample protein concentrations were measured by bicinchoninic acid (BCA) assay (23225; Thermo Scientific). Samples were then reduced with 10 mM dithiothreitol (DTT) at 95 °C for 5 min, cooled, then alkylated with 30 mM iodoacetamide (IA) for 30 min in the dark without shaking at room temperature.

Subsequently, 10 mM DTT was added to each sample, followed by a quick vortex, and incubation for 10 min at room temperature without shaking. Sample digestion was performed using sequencing grade trypsin (V5113; Promega) overnight at 37 °C, with generated peptide pools quantified by Nanodrop, normalized to 400 µg and acidified with formic acid (FA) to a final concentration of 5% (v/v) prior to being desalted using 1cc tC18 Sep Pak cartridges (WAT036820; Waters) as previously described^82^.

### 2.4 LC-MS/MS analysis

Changes in the protein abundance were assessed using trypsin digested samples analysed using a Fusion Lumos Tribrid Orbitrap mass spectrometer (Thermo Scientific) in a data independent acquisition (DIA) mode using the BoxCarDIA method^76,77^. Dissolved peptides (1 µg) were injected using an Easy-nLC 1200 system (LC140; Thermo Scientific) and separated on a 50 cm Easy-Spray PepMap C18 Column (ES803A; Thermo Scientific). A spray voltage of 2.2 kV, funnel RF level of 40 and heated capillary at 300 °C was deployed, with all data acquired in profile mode using positive polarity with peptide match off and isotope exclusion selected. All gradients were run at 300 nL/min with analytical column temperature set to 50 °C. Peptides were eluted using a segmented solvent B gradient of 0.1% (v/v) FA in 80% (v/v) ACN from 4% - 41% B (0 - 107 min). BoxCar DIA acquisition was as previously described^76^. MS1 analysis was performed by using two multiplexed targeted SIM scans of 10 BoxCar windows each, with detection performed at a resolution of 120,000 at 200 m/z and normalized AGC targets of 100% per BoxCar isolation window. Windows were custom designed as previously described^76^. An AGC target value for MS2 fragment spectra was set to 2000%. Twenty-eight 38.5 m/z windows were used with an overlap of 1 m/z. Resolution was set to 30,000 using a dynamic maximum injection time and a minimum number of desired points across each peak set to 6.

### 2.5 Data Analysis

All acquired BoxCar DIA data was analyzed in a library-free DIA approach using Spectronaut v14 (Biognosys AG) default settings. Key search parameters employed include: a protein, peptide and PSM FDR of 1%, trypsin digestion with 1 missed cleavage, fixed modification including carbamidomethylation of cysteine residues and variable modifications including methionine oxidation. Data was Log2 transformed, globally normalized by median subtraction with significantly changing differentially abundant proteins determined and corrected for multiple comparisons (Bonferroni-corrected p-value ≤ 0.05; q-value <0.05).

### 2.6 Bioinformatics

Retrieval for conversion of canola gene identifiers to Arabidopsis gene identifiers (AGI) for Gene Ontology enrichment, STRING association network analysis and SUBA5 subcellular localization was performed using Uniprot (https://www.uniprot.org/). Gene Ontology (GO) enrichment analysis was performed using The Ontologizer (http://ontologizer.de/;) to perform a parent-child enrichment analysis.

All proteins quantified within the study served as the background and significance was determined using the p-value <0.01. GO dot plots were made using *R* version 4.3.1 and R package *ggplot2.* Predicted subcellular localization information was obtained using SUBA5 and the consensus subcellular localization predictor SUBAcon (https://suba.live/). STRING association network analysis was performed in Cytoscape (version 3.10.1; https://cytoscape.org/) using the String DB plugin (https://string-db.org/) and enhancedGraphics (version 1.5.5; https://apps.cytoscape.org/apps/) Cytoscape applications. The minimum edge threshold was set to 0.9 for the shoot network and 0.7 for root. All data were plotted using GraphPad Prism (version 8; https://www.graphpad.com/scientific-software/prism/). Final figures were assembled using Affinity Designer software (version 2.4.1; https://affinity.serif.com/en-us/designer/).

### 2.7 Arabidopsis root growth analysis

*Arabidopsis thaliana* ecotype Col-0 was used as wild-type and T-DNA insertion lines *hip1* (SALK_059751C), *cml42* (SALK_041400C), *iqd32* (SALK_108620C), and *chmp1b* (SALK_135944C) were obtained from the Arabidopsis Biological Resource Centre (ABRC) and genotyped to homozygosity. After sterilization as previously described above for canola seeds, T-DNA line seeds with Col-0 control were germinated on media plates containing ½ MS either complete (Caisson Labs, MSP01), or nutrient deficient plates: -N (Phytotechnology Laboratories, M531), -P (Caisson Labs, MSP11), or -S (Caisson Labs, MSP44) with 7 g/L agar (Sigma) at pH 5.8. The media plates were placed vertically in a black 3D printed stand to limit root light exposure and germinated at a 12 h light at 21 °C and 12 h dark at 19 °C photoperiod. Primary root length measurements were taken over a 10-day time course, with representative photographs obtained on the 10^th^ day.

## 3 RESULTS

### 3.1 Plant growth in nutrient deficient conditions

To analyze the effects of nutrient deficiency on canola, we grew Westar canola plants in PRO-MIX® BY MYCORRHIZAE™ mixture with no additionally applied nutrients for 14 days. Next, the plants were fertilized with four different versions of Hoagland’s solution, resulting in treatments deficient in nitrogen (-N), phosphorus (-P), potassium (-K), and sulphur (-S) (**Figure 1a**). Plants were then grown for a further 10 days prior to sampling of both shoots and roots. We first verified that the plants were indeed experiencing nutrient deficiency within our experimental system by performing inductively coupled plasma - optical emission spectrometry (ICP-OES) analysis to quantify the levels of N, P, K and S in shoot tissues obtained for each experimental condition. As shown in **Figure 1b**, we detected significant reductions in the desired nutrient deficiency for each treatment. Interestingly, we also observed a significant reduction in K under the -N treatment, despite this sample treatment having a K concentration consistent across all nutrient deprivation conditions, suggesting a possible role for N in K uptake and/or transport. Additionally, although not significant, we also observed a decreased amount of shoot K under the -P condition (**Figure 1b**). This is consistent with a previous report in *Glycine max*, which found K concentrations in plant tissues reduced by half under P deficient conditions^83^.

**Figure 1.**
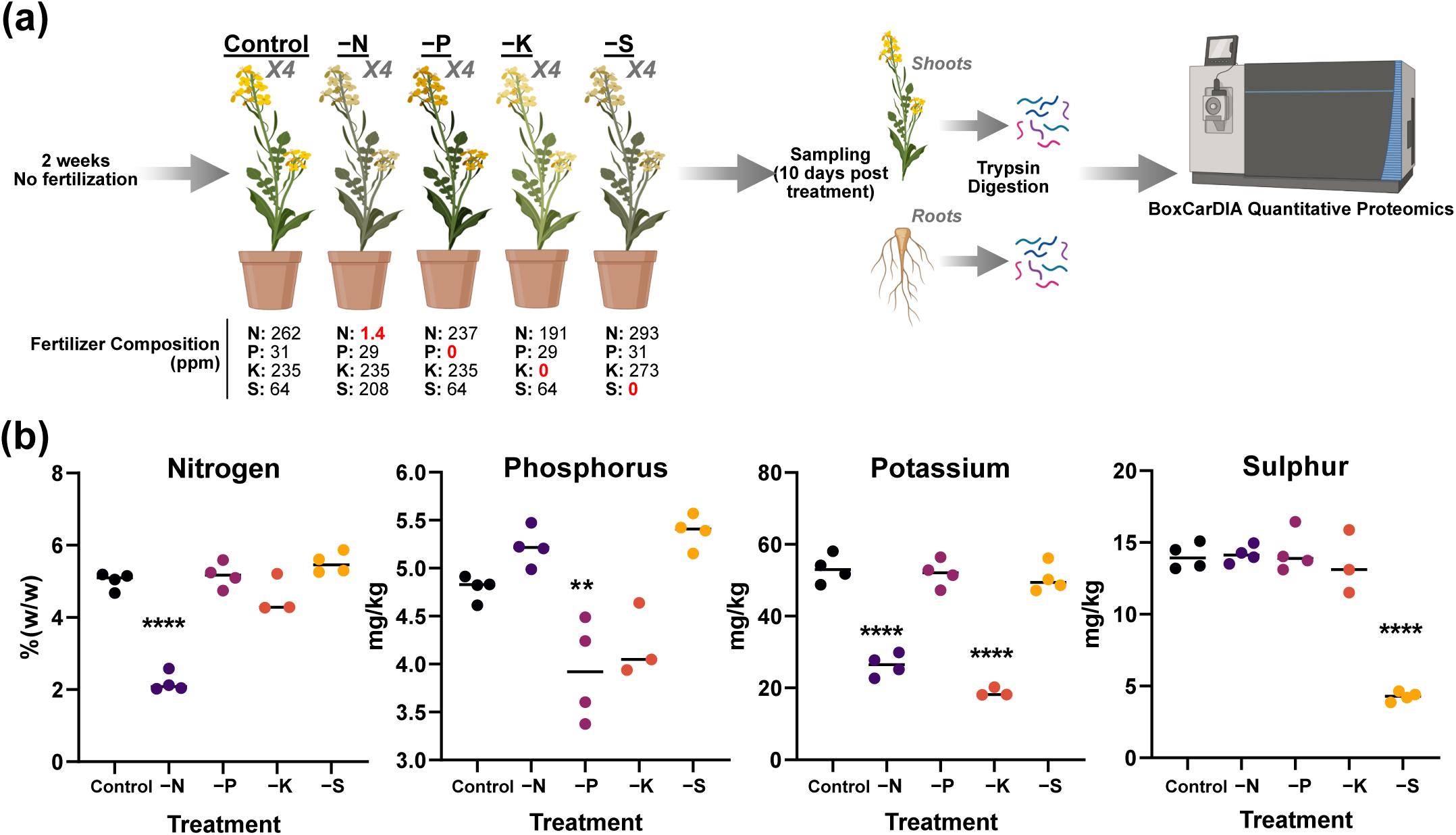
Experimental design and validation of nutrient deficiency a) Canola were grown for 14 days with no fertilization followed by growth under Control, -nitrogen (-N), -phosphorus (-P), -potassium (-K), or -sulphur (-S) conditions for 10 days. Shoots and roots were separated upon harvesting prior to protein extraction, trypsin digestion and BoxCarDIA quantitative proteomic analysis. b) ICP-OES testing for total nitrogen (%w/w), total phosphorus, total potassium, and total sulphur (mg/kg) in canola shoot tissue under the experimental Control, -N, -P, -K, and -S treatments as described in (A). Statistical test used was a one-way ANOVA comparing control and each treatment, followed by Dunnett’s multiple comparisons test. Stars denote treatments that were significantly different from control. **, p-value <0.01; ****, p-value <0.0001.

We next employed the use of high-resolution LC-MS/MS to quantify changes in the shoot and root proteome across these different nutrient deficient conditions. Overall, we were able to quantify over 4,000 proteins across all the shoot samples, and over 1,100 proteins across all the root samples (**Table 1; Supp Table 2**). We next performed fold-change analysis for each of our treatment conditions compared to the control, resolving several hundred significantly changing proteins for each treatment and tissue type (Log2FC > 0.58; q-value < 0.05) (**Table 1**). Interestingly, a third of the proteins quantified in our -N shoot samples displayed significant changes in abundance. This was also far greater than the ∼20% of proteins that showed significant changes in the -N root samples. Additionally, while we quantified overall fewer proteins in roots compared to shoots in each of our nutrient deficient treatments, a higher percentage of root proteins exhibited abundance changes compared to the number of shoot proteins.

**Table 2.** Average Log2FC values for significantly changing proteins that were differentially regulated in shoot vs. root under at least one nutrient stress.

| Brassica ID | AGI | Gene Name | Average log2FC |  |  |  |  |  |  |  |
| --- | --- | --- | --- | --- | --- | --- | --- | --- | --- | --- |
|  |  |  | Shoot -N | Shoot -P | Shoot -K | Shoot -S | Root -N | Root -P | Root -K | Root -S |
| BnaA01T0044600WE | AT4G22670 | HIP1 | -2.12726 | 0 | -1.9714 | 0 | 0 | 0.952935 | 1.411874 | 0 |
| BnaA08T0110100WE | AT4G22670 | HIP1 | -2.32124 | -1.3544 | 0 | -1.243404 | 0.811867 | 1.308123 | 1.359938 | 0.815155 |
| BnaA10T0292900WE | AT5G02070 |  | -1.80422 | 0 | -0.62613 | -0.606195 | 0.831729 | 1.088581 | 1.156854 | 0 |
| BnaC09T0279900WE | AT4G10480 |  | -1.69888 | 0 | -1.16478 | 0 | 1.351642 | 1.095766 | 0.766503 | 0 |
| BnaC01T0106600WE | AT4G20780 | CML42 | -1.59409 | -1.20071 | -1.30729 | -1.153585 | 2.480891 | 3.234212 | 1.254506 | 2.630429 |
| BnaA09T0010800WE | AT4G00830 | LIF2 | -1.37899 | 0 | 0 | 0 | 0.860354 | 0.701917 | 0 | 0 |
| BnaA01T0282800WE | AT3G12390 |  | -1.31614 | 0 | 0 | 0 | 0.738326 | 0 | 0.728514 | 0 |
| BnaA03T0394200WE | AT3G25230 | FKBP62 | -1.19984 | 0 | 0 | 0 | 0.778684 | 1.549793 | 1.147278 | 0 |
| BnaA06T0301500WE | AT4G26110 | NAP1;1 | -1.17582 | 0 | 0 | 0 | 1.42574 | 0.961554 | 0 | 0 |
| BnaA06T0413800WE | AT3G27530 | GC6 | -1.12537 | 0 | 0 | -0.619403 | 1.065348 | 0.670267 | 0 | 0 |
| BnaC09T0245200WE | AT2G04030 | HSP90.5 | -0.89609 | 0 | 0 | 0 | 1.033867 | 0 | 0 | 0 |
| BnaA09T0026100WE | AT4G03520 | TRXM2 | -0.6695 | 0 | 0 | 0 | 1.130324 | 1.230541 | 1.22092 | 0 |
| BnaA01T0284400WE | AT3G11940 | RPS5A | -0.62721 | 0 | 0 | 0 | 2.574966 | -1.27309 | -2.45336 | 0 |
| BnaA02T0201400WE | AT1G17730 | CHMP1B | -0.58162 | -0.62319 | 0 | 0 | 2.664905 | 2.19228 | 2.475491 | 2.052978 |
| BnaC08T0188800WE | AT1G15340 | MBD10 | 0.585919 | -0.74398 | -1.1754 | 0 | -0.81628 | 0 | 0 | -1.00673 |
| BnaC04T0574100WE | AT2G40300 | FER4 | 0.590266 | 0 | -1.53158 | 0 | -0.75866 | 0 | 0 | 0 |
| BnaC09T0465700WE | AT5G17380 | HPCL | 0.610642 | 0 | 0 | 0 | -5.13574 | 0 | 0 | -3.63132 |
| BnaA03T0361400WE | AT3G17390 | MAT4 | 0.668672 | 0 | 0.729214 | 0 | -1.67312 | 0 | 0 | 0 |
| BnaA10T0108600WE | AT5G55070 | E2-OGDH2 | 0.759383 | 0 | 0 | 0 | -1.23714 | -2.24159 | 0 | -1.18072 |
| BnaA04T0034000WE | AT3G56240 | HMP31 | 0.840463 | 0 | 0 | 0 | -1.24129 | 0 | 0.842226 | 0 |
| BnaA03T0011500WE | AT5G01600 | FER1 | 1.212664 | 0.636781 | -1.28568 | 0 | -1.40946 | 0 | 0 | 0 |
| BnaA10T0238900WE | AT5G10270 | CDKC;1 | 1.626956 | 0 | -1.38006 | 0 | -1.55804 | 0.89802 | 0 | 0 |
| BnaC05T0297800WE | AT1G48620 | HON5 | 2.240977 | 0 | -1.44833 | 1.712588 | -1.90114 | -0.68174 | 0 | 0 |
| BnaC05T0417600WE | AT3G15950 | NAI2 | -2.00872 | -1.27397 | -1.51455 | 0 | 0 | 0.991 | 0 | 0 |
| BnaA10T0058300WE | AT1G47710 | SERPIN1 | 0 | -0.89739 | 0 | -0.938194 | 1.023878 | 0.721894 | 0.812397 | 0 |
| BnaC02T0243900WE | AT1G70830 | MLP28 | 0 | -0.82983 | -0.9176 | 0 | 0 | 0.920672 | 1.19166 | 0.949179 |
| BnaA07T0239700WE | AT1G75950 | SKP1 | 0 | -0.7026 | -0.71413 | 0 | 0.941403 | 1.044871 | 1.352355 | 1.059284 |
| BnaC01T0329400WE | AT3G19390 |  | 0 | -0.66264 | 0 | 0 | 1.789872 | 2.618133 | 1.929591 | 1.654872 |
| BnaA04T0168300WE | AT2G27510 | FD3 | 0 | -0.65528 | 0 | -0.74641 | 0 | 0.664527 | -0.84829 | 0 |
| BnaA04T0016500WE | AT3G60240 | EIF4G | 0 | -2.54719 | -3.21005 | -0.789307 | 0 | 0 | 0.872639 | 0 |
| BnaA09T0548600WE | AT2G24420 |  | 0.889844 | -1.17179 | -2.5337 | -0.59908 | 0 | 0 | 0.741261 | 0 |
| BnaC04T0086100WE | AT2G38040 | CAC3 | 0 | -1.33175 | -1.72718 | -0.677862 | 0 | 0 | 1.111734 | 0 |
| BnaC05T0246500WE | AT2G18110 |  | -1.24798 | -1.55839 | -1.46916 | -1.482333 | 0 | 0 | 1.857518 | 0 |
| BnaC04T0467900WE | AT2G27710 | P2Y | -1.18515 | -1.12317 | -1.46674 | 0 | 0 | 0 | 1.170042 | 0.610114 |
| BnaC01T0162700WE | AT4G27580 | P2Y | 0 | -0.94357 | -1.33004 | -0.628645 | 0 | 0 | 1.775323 | 0 |
| BnaA07T0105900WE | AT1G26665 | MED10B | -1.32801 | 0 | -1.03501 | 0 | 0 | 0.862857 | 1.416753 | 0 |
| BnaA02T0064600WE | AT5G16590 | QSK2 | 0 | -1.1426 | -0.92546 | -1.301945 | 0 | 0 | 2.493405 | 0 |
| BnaC06T0125700WE | AT1G55670 | PSAG | 0 | 0 | -0.877 | 0 | 1.055415 | 1.022279 | 0.615953 | 0 |
| BnaA10T0225700WE | AT5G12140 | CYS1 | -0.83715 | 0 | -0.81765 | 0 | 0 | 2.730179 | 1.625106 | 1.594633 |
| BnaA06T0097500WE | AT1G14980 | CPN10 | 0 | -0.59222 | -0.77752 | 0 | 0 | 0 | 0.784039 | 0 |
| BnaC01T0077300WE | AT4G17520 | HLN | 0 | 0 | -0.70176 | 0 | 0 | 0.741798 | 0.857577 | 0 |
| BnaC07T0223800WE | AT5G44130 | FLA13 | 0 | 0 | -0.69941 | 0 | 0 | 1.072278 | 0.777888 | 1.028012 |
| BnaA03T0400100WE | AT2G06530 | VPS2.1 | 0 | 0 | -0.68199 | 0 | 0 | 1.296335 | 1.474631 | 0 |
| BnaA08T0117800WE | AT4G34450 | GAMMA2-COP | 0 | -1.20064 | -0.65725 | 0 | 0 | 0 | 1.340956 | 0 |
| BnaA07T0149800WE | AT2G27730 | P2 | 0 | 0 | 0.679475 | 0.696413 | 0 | 0 | -8.85036 | 0 |
| BnaA07T0227200WE | AT1G78370 | GSTU20 | -0.76773 | 0.581809 | 0.740131 | 0 | -1.81833 | 0 | -2.16382 | 0 |
| BnaC03T0108100WE | AT2G30060 |  | 0 | 0.857036 | 1.002155 | 11.13647 | 0 | 0 | -2.72422 | 0 |
| BnaA02T0252900WE | AT4G04720 | CPK21 | -2.58494 | 0 | -2.39778 | -1.288472 | -0.8626 | 1.889768 | 0 | 0.722297 |
| BnaA09T0200400WE | AT5G47210 |  | 0 | 0 | -1.49484 | 0.73302 | 0 | 0 | 0 | -0.69033 |

### 3.2 Correlation analysis identifies relationships between changing proteins and nutrient stressors

To assess proteome dynamics in both shoots and roots under the different nutrient deficiency conditions, we next performed a correlation analysis comparing the proteome fold-changes observed in each stress conditions for each organ (**Figure 2a, b**). This revealed striking correlations in proteome-level changes between different pairs of nutrient deficiencies in shoots and roots. For instance, when comparing shoot and root proteomes, we found that root proteomes across nutrient deprivation conditions responded more similarly to each other than did shoots. Additionally, this analysis revealed a strong correlation between the -P and -S treatments in both shoot and root proteomes, while -N and -S resulted in a more divergent correlation in shoots, but similar changes in roots. To better contextualize these correlations, we next performed Gene Ontology (GO) analysis of these correlated sub-populations (**Supp Table 3**). As *Arabidopsis thaliana* (Arabidopsis) has more tools to biologically contextualize our findings, we first identified the Arabidopsis orthologs for all significantly changing proteins from the *B. napus* genome annotation. We identified orthologous Arabidopsis gene identifiers (AGIs) for approximately 75.3% (2056/2731) of the quantified significantly changing canola protein population across the study. When separated into shoot and root proteomes, this amounted to 76.1% (1675/2200) and 72.1% (514/713) of significantly changing shoot and root proteomes, respectively.

**Figure 2.**
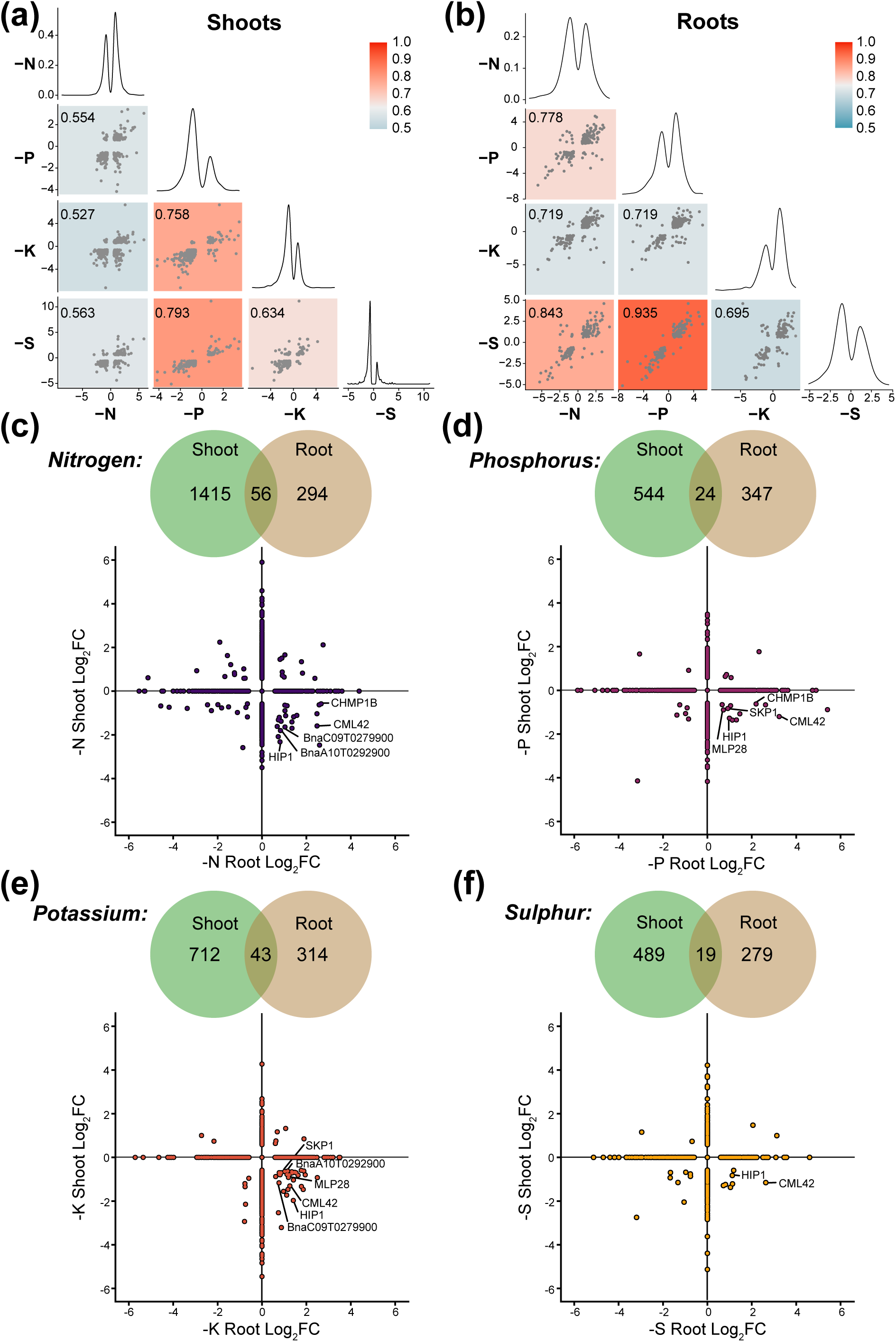
Correlation and distribution of significantly changing proteins in root and shoot under nutrient deficient conditions a-b) Correlation analysis of distribution of significantly changing proteins comparing -N, -P, -K and -S with each other in shoots (a) and roots (b). c-f) Venn diagrams of overlap of significantly changing proteins (q-value <0.05) common between or independent in roots vs. shoots under N (c), P (d), K (e), and S (f) nutrient deficiency. Distribution plots for each nutrient deficiency are shown below for significantly changing proteins common between root and shoot.

Enriched ‘Biological Process’ (BP)-GO terms for significantly changing shoot proteins exhibiting correlated changes in abundance in roots under -P and -S conditions include: ‘Chlorophyll Biosynthetic Process’ (GO:0015995) and ‘Positive Regulation of Tetrapyrrole Metabolic Process’ (GO:1901403), both of which largely consisted of downregulated proteins. Additionally, a number of BP-GO terms were also resolved in the root, including: ‘Negative Regulation of Endopeptidase Activity’ (GO:0010951), and ‘Negative Regulation of Proteolysis’ (GO:0045861), as well as ‘Cell Cycle’ (GO:0007049), and ‘DNA-Templated Transcription, Initiation’ (GO:0006352), which consisted mostly of upregulated proteins.

There was also an enrichment of BP-GO terms associated with general stress response such as ‘Response to Osmotic Stress’ (GO:0006970), ‘Protein Folding’ (GO:0006457), and ‘Response to Endoplasmic Reticulum Stress’ (GO:0034976), also consisting of proteins that were upregulated under the nutrient deficiency. In the root, several ‘Molecular Function’ (MF)-GO terms showed an enrichment under -P and -S conditions comprised of proteins involved in ‘Calcium Ion Binding’ (GO:0005509), ‘Oxidoreductase Activity’ (GO:0016684), and ‘Endopeptidase Inhibitor Activity’ (GO:0004866), each of which encompassed significantly upregulated proteins (**Supp Table 3**).

Additionally, we find different enriched BP-GO and MF-GO terms for significantly changing proteins exhibiting correlated changes in their abundance between -N and -S growth conditions in the root. These include: ‘Cellular Amino Acid Catabolic Process’ (GO:0009063), and ‘Alpha-Amino Acid Catabolic Process’ (GO:1901606), which included proteins involved in glycine decarboxylation.

Similarly to the significantly changing, correlated -P and -S root proteins, we find an enrichment of proteins with the MF-GO term ‘Oxidoreductase Activity’ (GO:0016684), which is comprised of a mixture of up- and down-regulated proteins, and ‘Cysteine-Type Peptidase Activity’ (GO:0008234), with mostly upregulated proteins.

To determine whether there were distinct similarities or differences between the subcellular localization of the changing proteome within and between nutrient deficiency conditions, we performed subcellular localization analysis for shoots and roots under each experimental condition using SUBA consensus analysis tool (SUBAcon; https://suba.live/; **Figure 3**). Interestingly, shoots in general had more extensive proteome changes relating to proteins localized to the plastid and peroxisome localized proteins, while roots had far more proteome changes relating to nuclear proteins. In particular, -P conditions in the root had a disproportionate number of cytosolic proteins changing in their abundance compared to all other conditions.

**Figure 3.**
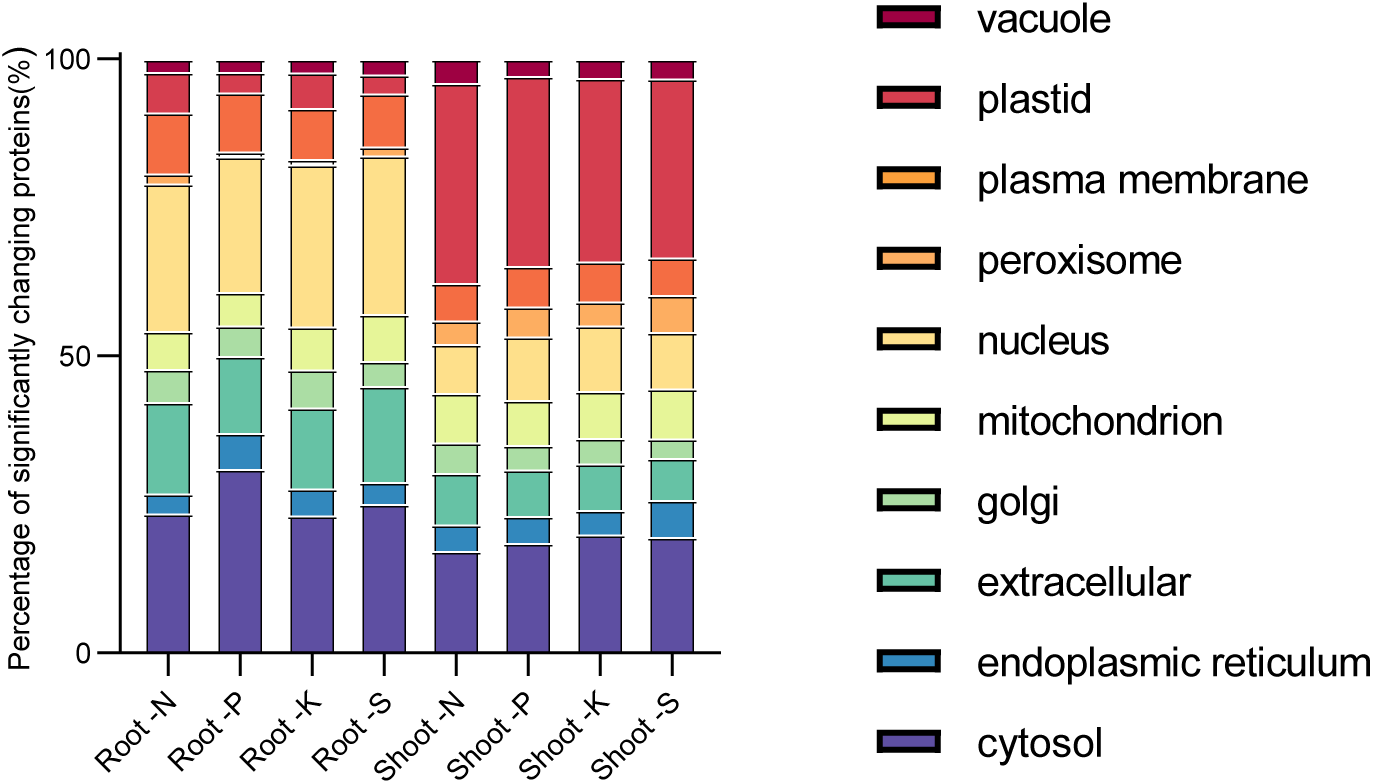
Analysis of subcellular localization *In silico* subcellular localization analysis of significantly changing proteins was performed using SUBAcon (https://suba.live/) using converted Arabidopsis gene identifiers.

Comparing shoot and root proteome changes in each nutrient deficiency condition revealed clear organ-specific partitioning in responses as only 56, 24, 43, and 19 canola proteins were found to change in both organs for -N, -P, -K, and -S, respectively (**Figure 2c-f**). Intriguingly, a sizeable number of these proteins displayed opposite changes in abundance between shoots and roots for each stress (**Table 2**). For example, the calmodulin-like protein, CML42 (BnaC01T0106600), and an HSP70-interacting protein, HIP1 (BnaA08T0110100; BnaA01T0044600) were downregulated in shoot and upregulated in root across all four nutrient deficiency conditions. The subunit of the ESCRTIII complex, CHMP1B

(BnaA02T0201400), was downregulated in shoot and upregulated in root in the -N and -P conditions. A receptor-like protein kinase, BnaA10T0292900, and a NAC family protein, BnaC09T0279900, were downregulated in shoot and upregulated in root in both the -N and -K conditions and MAJOR LATEX PROTEIN (MLP28; BnaC02T0243900) and S-PHASE KINASE ASSOCIATED PROTEIN (SKP1; BnaA07T0239700) were downregulated in shoot and upregulated in root in both the -P and -K conditions. Additionally, there were several other proteins exhibiting similar abundance change patterns in individual nutrient deficient conditions (**Table 2**). It is also striking that both -N and -K treatments result in a greater number of proteins upregulated in the shoot and downregulated in the root than the -P and -S conditions. Overall, this highlights the necessity of investigating proteome changes in different organs and across multiple related stress scenarios.

### 3.3 GO analysis of nutrient-deficiency induced shoot and root proteome changes reveals similarities between the nutrient stresses

To gain a more thorough understanding of the processes affected by either individual or multiple nutrient deficiency conditions in the shoot or root, we next performed GO enrichment analyses on significantly up- and down-regulated proteins (Log2FC >0.58, q-value <0.05) for each nutrient deficiency condition in shoot and root (**Figure 4; Supp Figure 1**). We identified several GO terms in both shoots and roots that were enriched across multiple nutrient stress treatments. In shoots, we observed a general enrichment of BP-GO terms involved in photosynthesis, and chlorophyll biosynthesis and metabolism amongst the population of downregulated proteins including ‘Chlorophyll Biosynthetic Process’ (GO:0015995) and ‘Tetrapyrrole Biosynthetic Process’ (GO:0033014) under -N, -P, and -S growth conditions, and ‘Photosynthesis’ (GO:0015979) under -N and -K growth conditions (**Figure 4a**). For upregulated shoot proteins under nutrient deficiencies, the enriched BP-GO terms primarily reflected changes in carbohydrate metabolism and starch breakdown, including the enrichment of BP-GO terms ‘Glucan Catabolic Process’ (GO:0009251) and ‘Polysaccharide Metabolic Process’ (GO:0005976) under the -N and -K conditions. Additionally, enriched GO terms for amino acid and N compound catabolism were observed, including ‘Alpha-Amino Acid Catabolic Process’ (GO:1901606) under -N and -P growth conditions, and ‘Organonitrogen Compound Catabolic Process’ (GO:1901565) under the -N, -P and -S treatments (**Figure 4a**). This indicates more complex C and N partitioning responses in aboveground tissues relative to belowground tissues as a response to different nutrient deficiencies.

**Figure 4.**
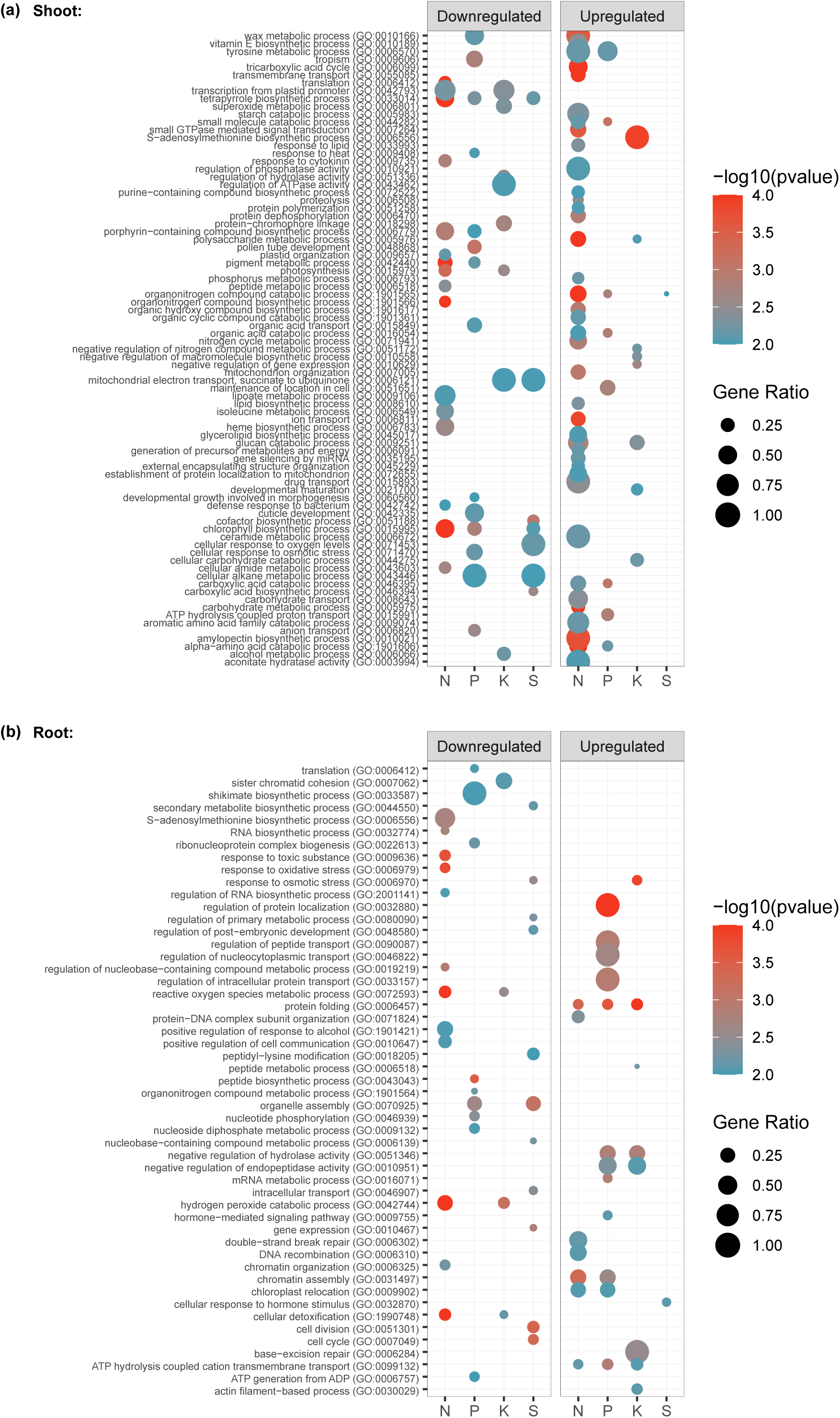
Gene Ontology (GO) enrichment analysis Dotplot representation of enriched biological process GO terms for significantly changing proteins (q- value <0.05; Log2FC >0.58 or <-0.58) that are up- or downregulated in shoot (A) or root (B) under each nutrient deficiency (-N, -P, -K or -S). The size of the dots represents the gene ratio (number of proteins in the conditions/total quantified proteins in the study). The colour code of the dots represents the log10(p- value). Only GO categories with p-value <0.01 and total count <500 are included.

In roots, BP-GO terms that were enriched for downregulated protein groups across multiple nutrient stress conditions largely consisted of those related to oxidative stress response, including ‘Cellular Detoxification’ (GO:1990748), ‘Hydrogen Peroxide Catabolic Process’ (GO:0042744), and ‘Reactive Oxygen Species Metabolic Process’ (GO:0072593), which were downregulated under -N and -K conditions (**Figure 4b**). Additionally, ‘Oxidoreductase Activity’ (GO:0016684) was enriched for downregulated proteins under -N and -K treatments. Hydrogen peroxide and ROS act as signaling molecules in root development and elongation^84^, which suggests that this downregulation may facilitate accumulation of ROS, accounting for the altered root system architecture observed under nutrient stress conditions.

In roots, proteins that were upregulated in their abundance under nutrient deficient conditions were enriched in BP-GO terms relating to ‘ATP Hydrolysis Coupled Cation Transmembrane Transport’ (GO:0099132) under -N, -P and -K conditions (**Figure 4b**), and MF-GO terms ‘Active Transmembrane Transporter Activity’ (GO:0022853) under -P and -K conditions (**Supp Figure 1b**), indicating increased active transport in roots under nutrient stress conditions. Additionally, there was an upregulation of proteins associated with endopeptidase activity and its regulation, with the enrichment of BP-GO terms including ‘Negative Regulation of Endopeptidase Activity’ (GO:0010951) and ‘Negative Regulation of Hydrolase Activity’ (GO:0051346) under -P and -K treatments, as well as enriched MF-GO terms ‘cysteine-type peptidase activity’ (GO:0008234) across all nutrient stress conditions, and ‘endopeptidase inhibitor activity’ (GO:0004866) and ‘hydrolase activity, acting on acid anhydrides, catalyzing transmembrane movement of substances’ (GO:0016820) under -P and -K only. Finally, in roots we find enriched GO terms for proteins increasing in abundance suggestive of general stress responses. These include: ‘Protein Folding’ (GO:0006457) under -N, -P, and -K conditions, and ‘Calcium Ion Binding’ (GO:0005509) under -P, -K and -S nutrient deficient conditions.

### 3.4 Network analysis of shoot and root proteome changes identify commonalities under nitrogen, phosphorus, potassium and sulphur deficiency stress

To further contextualize our GO enrichment analysis and identify clusters of associated proteins within and among the nutrient deficiencies in shoot and root, we performed a STRING-DB association network analysis for the shoot and root using the respective significantly changing protein populations (**Figure 5**). Relative to Arabidopsis, canola has undergone substantial genome changes, triplicating in size^85^. Thus, we observed several duplicate canola homologs accounting for one single Arabidopsis gene. Upon inspection, the duplicates did not consistently follow the same pattern of protein differential expression across nutrient deficiencies, and thus to prevent biasing the analysis, we identify the duplicates as a separate green colour on the STRING-DB association network (**Supp Table 4**).

**Figure 5.**
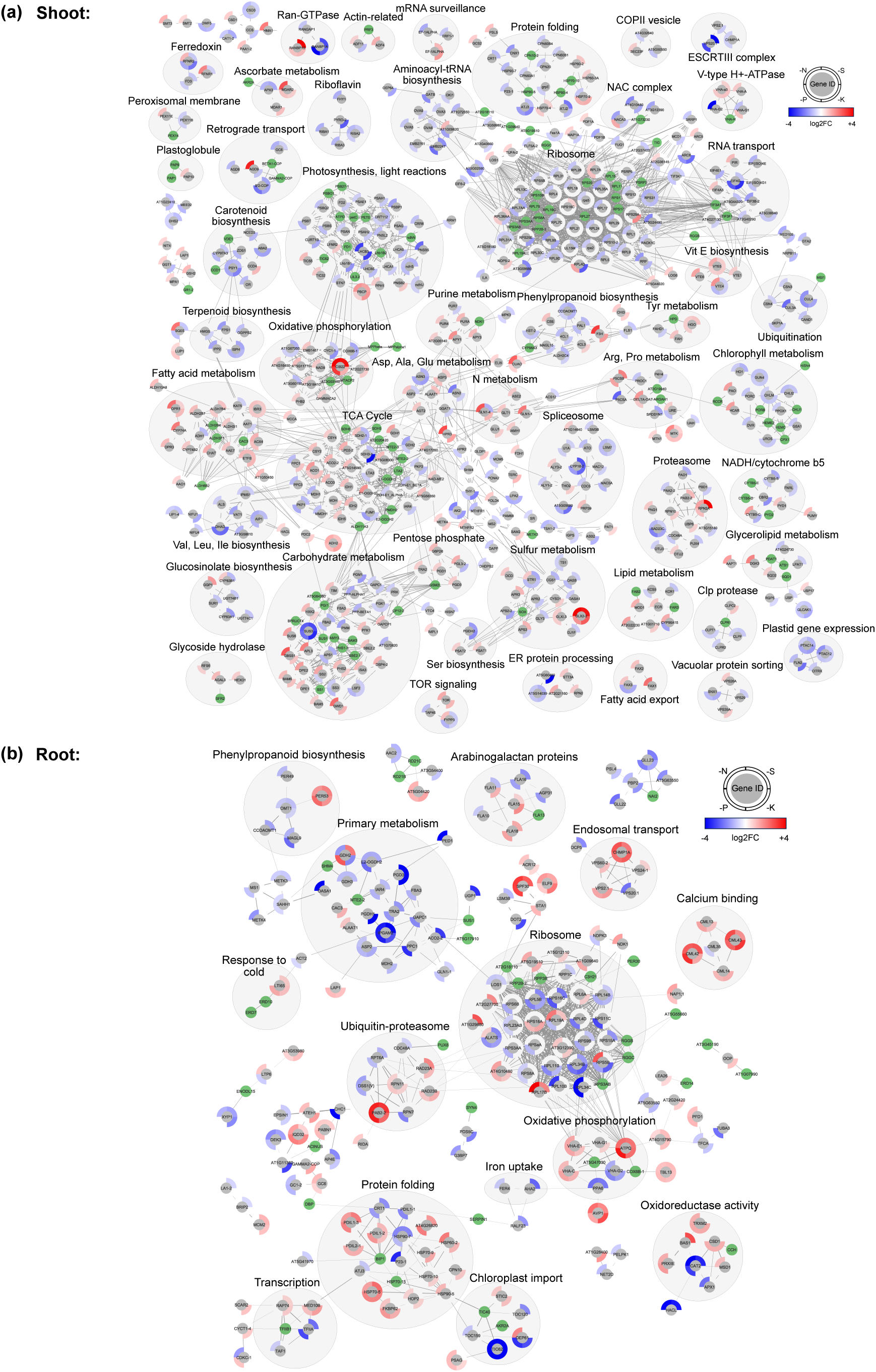
Association network analysis of proteins changing under nutrient deficiencies An association network analysis was performed using STRING-DB (ANOVA p-value <0.05) depicting significantly changing proteins under all nutrient deficiency treatments in shoots (a) and roots (b). Edge thickness indicates the strength of the connection between nodes. Minimum edge threshold was set to 0.9 for (a) and 0.7 for (b). Significantly changing proteins are labelled by either their primary gene annotation or Arabidopsis gene identifier (AGI). Outer circle around each node indicates the standardized relative Log2FC of the indicated protein in the nutrient deficiency vs. control. The scale of red to blue indicates relative increase and decrease in abundance, respectively. Groupings of nodes encompassed by a grey circle indicate proteins involved in the same cellular process.

In the shoot network, there were clear clusters showing downregulation of proteins involved in photosynthesis and light reactions, and chlorophyll metabolism across multiple nutrient deficiencies (**Figure 5a**), which we also observed in our shoot GO analysis (**Figure 4; Supp Figure 1**). We also uniquely observe differential regulation and a distinct upregulation of carbohydrate metabolic, TCA cycle and fatty acid metabolic proteins under -N conditions. Alternatively, in roots we observed clusters of proteins upregulated across multiple conditions for calcium binding and proteins folding responses (**Figure 5b**), validating the results of our root GO analysis (**Figure 4; Supp Figure 1**). We also observe clusters of downregulated proteins involved in C metabolism and phenylpropanoid metabolism. We further identify a cluster of proteins possessing oxidoreductase activity; however, it demonstrates more complex responses, with some proteins upregulated across multiple nutrient deficiency conditions and some downregulated. Collectively, when combined with GO analysis, our STRING-DB association network analysis offers a protein-level and process-level understanding of the nutrient-induced proteome changes experienced by canola shoots and roots.

### 3.5 Phenotypic validation nutrient deficiency-related putative candidates

In addition to the large number of proteins we observed changing in abundance that are known to be impacted by a deficiency in these nutrients, we see a number of unique candidates that seem to be affected under multiple nutrient stress conditions. Therefore, we next selected a series of novel nutrient-deficiency responsive candidates that we identified as significantly changing under all nutrient deficiency conditions in our study for further validation. To do this, we focused on assessing phenotypic changes in the roots, as they represent the primary interface of nutrient uptake. Specifically, we selected candidates that we found to be differentially abundant in shoots vs. roots under multiple conditions, totaling four candidate proteins. Homozygous T-DNA insertional knockout mutants were obtained for each candidate from the Arabidopsis Biological Resource Centre (ABRC; https://abrc.osu.edu/). Each line was germinated and grown on ½ MS media lacking individual nutrients for a -N, -P and -S treatment for 10 days. Of the four targets (*chmp1b, cml42, hip1,* and *iqd32*), we were able to validate the involvement of two candidates by assessment of primary root length changes under nutrient deficient conditions (**Figure 6; Supp Figure 2**). Specifically, under -N conditions *cml42* maintained a proportionally shorter primary root length on -N vs. control media (**Figure 6a-b**). Additionally, we find Col-0 plants have significantly longer primary roots under -S conditions than under control (**Figure 6c-d**). We did not observe this increased root length with the *iqd32* mutant, indicating a potential role in primary root elongation for sulphur scavenging response. Both IQD proteins and CMLs are involved in calcium binding, which we find enriched throughout our dataset, suggesting a broadly important role for calcium signaling in plant response to different nutrient deficiencies.

**Figure 6.**
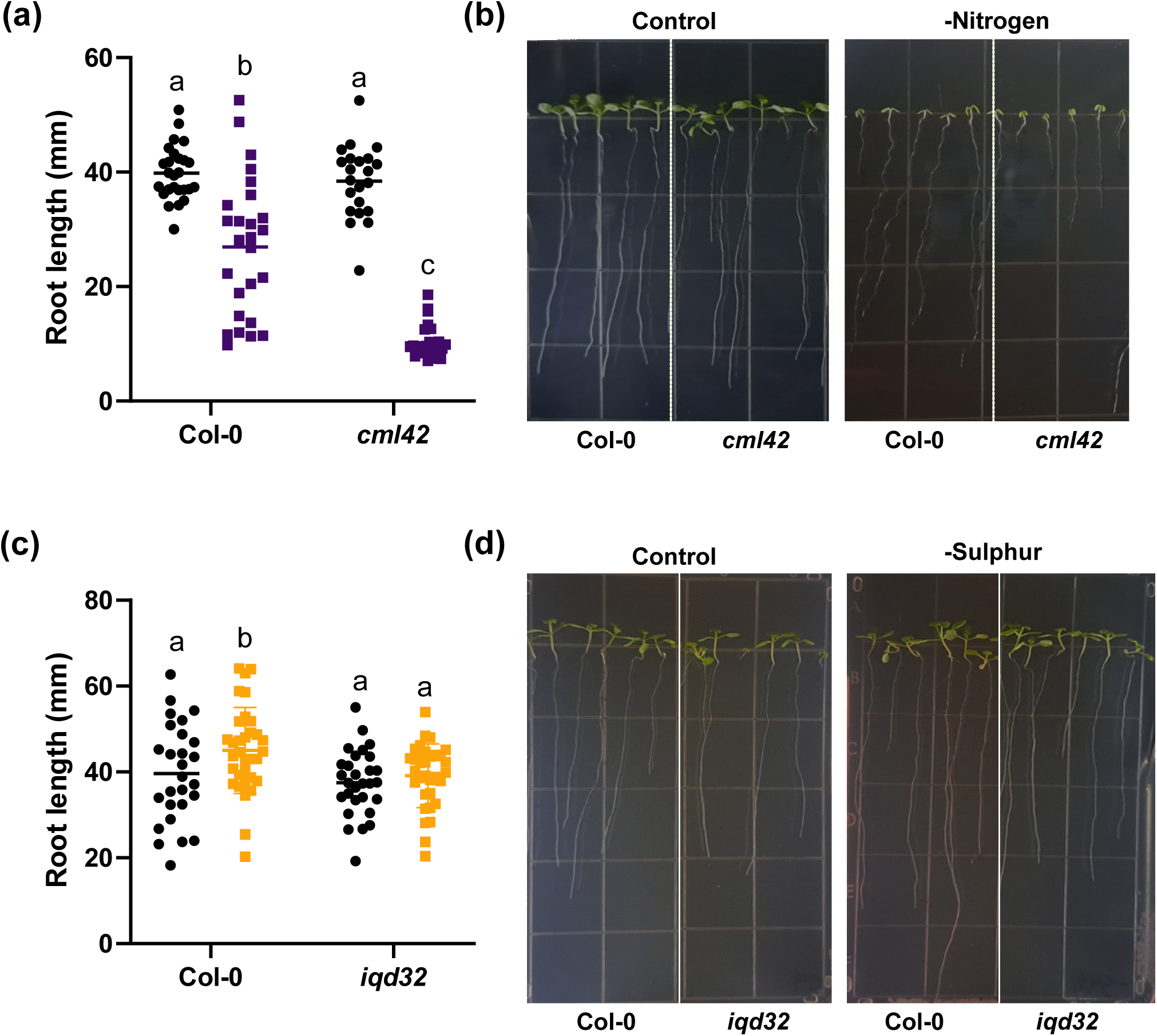
Root growth phenotypic validation of loss-of-function Arabidopsis seedlings grown under nutrient deficiency Root lengths were measured for wild-type (Col-0) and *cml42* or *iqd32* loss-of-function seedlings 10 days following germination on nitrogen-deficient (a-b) and sulphur deficient (c-d) media, respectively. A two- way ANOVA, followed by Fishers LSD test was used to determine statistical significance between nutrient-deficient conditions and genotypic background. Letters denote statistical significance (<0.05). *cml42*, calmodulin-like protein 42; *iqd32*, IQ-domain 32.

## DISCUSSION

### 4.1 Benchmarking proteome changes to hallmarks of nutrient deficiency response

We identified several changing proteins in our dataset to confirm that the nutrient treatments were sufficient to elicit a nutrient-deficiency response. Under -N, we observed an upregulation of many proteins involved in N transport and N assimilation and recycling through amino acid catabolism. The NITRATE TRANSPORTER/PEPTIDE TRANSPORTER FAMILY (NRT/NPF) proteins have fundamental roles in N uptake and transportation. We observed an upregulation of three NPF proteins in shoots tissues under -N conditions: NPF6.4/NRT1.3 (BnaA05T0293500), NPF5.10 (BnaC05T0189200), and NPF8.3 (BnaA06T0435300). In Arabidopsis, under -N conditions, *NRT2* genes have been found to account for 95% of high-affinity influx^86^. A study in canola found *NPF6.4* expression only induced by low N in roots^32^. Thus, no observable change NRT protein levels in roots, nor NRT2 protein-level changes in shoots was curious, emphasizing the importance of investigating both gene and protein expression. In addition, amino acids are a major N-source within plants, and their metabolism plays a fundamental role in N assimilation^87,88^. Accordingly, we observed changes in several proteins associated with amino acid metabolism. The glutamine synthetase/glutamate:2-oxoglutarate aminotransferase (GS/GOGAT) cycle is the primary mechanism of ammonia assimilation, with contributions from asparagine synthetase (ASN). In shoot under -N conditions, we observed upregulation of the GS; GLN1;1 (BnaA04T0088500) and GLN1;4 (BnaA10T0197000), confirming previous observations of transcriptional upregulation under low N in canola^89^. We also observe the upregulation of the canola GOGAT; GLT1 (BnaA03T0145200). Zhou et al.^89^ also found a decrease in the amount of asparagine under low N, consistent with our observation of downregulated ASN2 in shoot (BnaA06T0293800). This suggests canola may preferentially assimilate N into glutamine, rather than asparagine. Beyond protein-level changes in established -N responsive genes, we find several other proteins involved in amino acid metabolism were upregulated in shoots under -N conditions including: ORNITHINE-DELTA-AMINOTRANSFERASE (DELTA-OAT; BnaC07T0243600) and ARGINASE1 (ARGAH1;

BnaA03T0259800) involved in arginine catabolism, and AROMATIC ALDEHYDE SYNTHASE (AAS; BnaA07T0004000), FUMARYLACETOACETATE HYDROLASE (FAH; BnaA06T0083400), HOMOGENTISATE 1,2-DIOXYGENASE (HGO; BnaC02T0161300) and P-HYDROXYPYRUVATE DIOXYGENASE (PDS1; BnaA09T0641500), involved in phenylalanine and tyrosine catabolism. Plants also recycle N from pyrimidine nucleobases under N-starvation conditions^90,91^. Aligning with this, we observe upregulation of PYRIMIDINE1 (PYD1; BnaA01T0245400) and PYD2 (duplicates: BnaA10T0225300 and BnaC09T0507400) in shoots.

There are five families of inorganic phosphate (Pi) transporters in plants, PHOSPHATE TRANSPORTER (PHT)1-4 and PHOSPHATE1 (PHO1)^92^, of which we observe upregulation of PHO1;H3 (BnaC05T0108300) in the shoot under -P and -K conditions. No PHO1;H3 abundance changes were observed in roots. PHO1;H3 is recognized as a P-transporter controlling root-to-shoot P-translocation in Arabidopsis under zinc deficiency^93^, thus the abundance change observed in the shoot under -P conditions indicates additional systemic complexity. Another hallmark of P deficiency responses is the upregulation of purple acid phosphatases (PAPs)^94^. In shoot under -P conditions, we observed an upregulation of PAP1 (BnaA09T0605500) and PAP2 (BnaA08T0251200). Under P-deficiency, phospholipids are also replaced by sulfolipids through the activity of SULFOQUINOVOSYLDIACYLGLYCEROL (SQD) enzymes^95,96^. Correspondingly, we observed one of the three canola SQD1 orthologs upregulated in shoots (BnaC07T0453600). Interestingly, in roots we observed a downregulation of the PHOSPHOENOLPYRUVATE CARBOXYLASE (PEPC) isozyme PPC1 (BnaC01T0367000), which contrasts the marked transcriptional upregulation and increased PEPC activity observed in previous studies as well as its downregulation upon P-refeeding^97,98^. However, a more recent study in cotton also found a decrease in PEPC activity under low P growth conditions^99^. Lastly, we also saw a downregulation in roots of NON-SPECIFIC PHOSPHOLIPASE C3 (NPC3; BnaC05T0533000), a member of the non-specific phospholipase C protein family responsible for phospholipid replacement by galactolipids and sulfolipids under P-deficiency^100,101^.

In terms of K- and S-deficiency, there are significantly fewer known hallmark changes. Of those known, we observe an upregulation of the POTASSIUM TRANSPORTER/ K^+^ UPTAKE TRANSPORTER (POT4/KUP3; BnaA01T0328500), which was previously found to be upregulated under low K conditions^102^. While, under -S conditions, we find multiple proteins relating to S-containing amino acid metabolism and S assimilation to change in abundance. S is critical for the biosynthesis of amino acids cysteine and methionine, with inorganic S first fixed into cysteine, and then methionine is synthesized from cysteine and O-phosphohomoserine through a trans-sulphation reaction^103^. In shoots, we observed a downregulation of O-ACETYLSERINE SULFHYDRYLASE (OASB; BnaA05T0030400), involved in cysteine biosynthesis, and of CYSTATHIONINE GAMMA-SYNTHASE1 (CGS1; BnaC05T0551400) and METHIONINE SYNTHASE2 (MS2; BnaA03T0302400), involved in methionine biosynthesis, indicating a decrease in S-containing amino acid biosynthesis. During S assimilation, following S uptake by transporters, S becomes activated to adenosine-5-phosphate (APS) by ATP sulfurylase (APS), and then reduced to sulfite by APS reductase (APR). Accordingly, we observed an upregulation in shoot of canola APS3 (BnaC01T0201200) and APR3 (BnaA03T0470200), indicating increased S-assimilation in the S-starved plants.

Taken together, we have found several protein-level changes related to genes previously established as responsive to each respective nutrient deficiency, validating our experimental approach. Interestingly however, many of these proteins do not strictly parallel previously observed transcriptional changes in canola, emphasizing the importance of quantifying protein-level changes in abundance.

### 4.2 Nutrient deficiency has consequences for photosynthesis and chlorophyll metabolism

By examining the effects of nutrient deprivation across four macronutrients in one study, we can reveal both common and unique responses that may inform future breeding practices. Common to all nutrient deficiency treatments, we discovered a general downregulation of proteins involved in photosynthesis, light reactions, and chlorophyll metabolism in shoots. Photosynthesis is essential for providing energy to support plant growth and development. There are many factors that contribute to photosynthetic efficiency, such as photosynthetic pigments (such as chlorophyll), photosystems, light-harvesting complexes and enzymes involved in C metabolism^104^, making it a complex energetic process that needs to be tightly regulated. Many cellular processes involved in energy generation and growth require essential macronutrients N, P, K, and S. For instance, N impacts chlorophyll and light harvesting complex including photosystems (PS) I and II, oxygen evolving complex (OEC), cytochrome b_6_f complex, plastocyanin, ferredoxin, ferredoxin NADP^+^ reductase and ATP synthase^105^. P makes up ATP, NADPH, nucleic acids, sugar phosphates and phospholipids, which all play important roles in photosynthetic processes^106^. K has been shown to promote chlorophyll biosynthesis^107^, and although it is not a constituent of organic compounds in the way N and P are, its ionic form functions to activate enzymes involved in carbohydrate metabolism, and influences ATPase activity^108^. Finally, S is essential for Fe-S cluster formation in photosynthetic apparatus and electron transport chain^109^. Thus, plants under these nutrient deficiencies typically show depressed photosynthesis^99,104,106,110–112^. In our dataset, we observed a general downregulation of proteins corresponding to chlorophyll biosynthesis, pigment biosynthesis (especially carotenoids), photosynthesis, and light reactions in shoots under all nutrient deficiencies, but in particular for canola under -N and -K growth conditions. This included pigment-binding proteins, LIGHT HARVESTING COMPLEX1 (LHCA1; BnaC06T0226800), and LHCA5 (BnaC08T0034300), LHCB3 (BnaC09T0342700), and LHCB5 (BnaC03T0236900), NADPH dehydrogenase subunits PHOTOSYNTHETIC NADPH DEHYDROGENASE SUBCOMPLEX5 (PNSB5; BnaA02T0281400), PNSL2 (BnaA06T0090200), ndhN (duplicates: BnaC03T0065900; BnaA03T0123000), ndhS (BnaA01T0056700), and ndhU (BnaA10T0164800) and several others (**Figure 5**). Additionally, we observed a downregulation of proteins related to chlorophyll metabolism across all nutrient deficient conditions, including the chloroplastic-like magnesium chelatase subunit CHLI1 (duplicates: BnaA01T0014700; BnaC06T0182600), CHLI2 (BnaA02T0292700), which are key proteins involved in the insertion of magnesium into the chlorophyll molecule^113^, and PROTOCHLOROPHYLLIDE OXIDOREDUCTASE B (PORB; duplicates: BnaA01T0088100; BnaC07T0419600), and PORC (BnaA08T0297300), which are involved in the light-dependent reduction of protochlorophyllide a to chlorophyllide a, an essential step in chlorophyll biosynthesis^114^. In accordance with decreased chlorophyll biosynthesis, we observed an upregulation in proteins associated with chlorophyll catabolism.

A key step in chlorophyll breakdown involves 7-HYDROXYMETHYL CHL A REDUCTASE (HCAR)^115^, PHEOPHORBIDE A OXYGENASE (PAO), and RED CHLOROPHYLL CATABOLITE REDUCTASE (RCCR)^116^. In our dataset, we observed an upregulation of HCAR (BnaA10T0029700) and PAO (BnaA01T0161200) in -N conditions, and RCCR (duplicates: Bnascaffold286T0016300; Bnascaffold1716T0000500) under -N or -P conditions. Chlorophyll content largely dictates plant photosynthetic capacity^117^, thus this increase in proteins associated with chlorophyll catabolism suggests points of regulation for the reduction in the plant photosynthetic capacity required to cope with nutrient stress as well as potential recovery of N from chlorophyll compounds.

Lastly, during acclimation to light stress, state transitions adjust how light energy absorption is distributed between photosystem I and II^118^. We also observed a downregulation of the protein kinase STATE TRANSITION 7 (STN7; BnaC06T0374100) under -N conditions, and an upregulation of protein phosphatases PROTEIN PHOSPHATASE 1 (PPH1; BnaA03T0506100) under -N conditions, and of PHOTOSYSTEM II CORE PHOSPHATASE (PBCP; BnaA05T0133400) under all nutrient deficiency conditions; both of which regulate state transitions under plant light acclimation^119,120^. Taken together, our results indicate that under low nutrient stress, it is likely that plants are augmenting and/or decreasing their light absorption by altering state transitions to favour a lower-energy state for energy conservation.

### 4.3 Starch catabolism, TCA cycle and lipid metabolism proteins are upregulated under N deficiency

Our analysis found the greatest number of differentially regulated proteins in the shoots of -N canola plants. Approximately 74% (1235/1676) of these proteins were upregulated and involved carbohydrate metabolism, TCA cycle, or lipid/fatty acid metabolism. Closer inspection of proteins involved in carbohydrate metabolism revealed involvement in starch degradation. Consistent with these observations, N deficient leaves have been shown to possess decreased starch and an increased sucrose/starch ratio in *C. sinensis*^121^. Further, starch degradation genes were previously identified as upregulated in canola under N deficiency^122^. Interestingly, several of these starch degradation proteins were also upregulated under K deficiency, suggesting an intersection between N and K deficient carbohydrate metabolism responses. An upregulation of sugars under N deficiency has been determined by metabolite analysis in barley^123^, and high sucrose can negatively feedback on photosynthesis^124,125^, consistent with our observations of decreased photosynthetic protein abundance.

We also observed shoot-specific TCA cycle protein upregulation under -N conditions. Here, we find nearly all TCA cycle enzymes to be upregulated, except malate dehydrogenase (MDH) and pyruvate dehydrogenase (PDH), which are downregulated. A similar phenomenon was reported in rice leaves^126^. These results suggested an overall plant strategy of reduced TCA cycle flux under -N conditions by reducing pyruvate conversion to acetyl CoA, while increasing malate levels, which could provide carbon to aid in N assimilation^127^. Thus, a decrease in PDH and MDH levels might serve to regulate the amount of N going into amino acid biosynthesis.

Lastly, we also find fatty acid metabolic proteins upregulated under canola -N conditions. The majority of these significantly upregulated proteins correspond to the biosynthesis or elongation of very long chain fatty acids and waxes. Previous studies have shown changes in galactolipid and wax compositions in Arabidopsis and rice, respectively under -N conditions^128,129^. In accordance with our observations, there is also a report in maize of decrease in medium chain fatty acids but increase in very long chain fatty acids in low N conditions^130^, which form waxes and could be related to cell wall alterations. Interestingly, to date almost nothing is known about the role of very long chain fatty acids in nutrient stress responses, offering multiple interesting avenues for further investigation.

### 4.4 Nutrient deficiency induces downregulation of oxidative stress responsive proteins in canola roots

In our dataset, we observed the highest correlation between nutrient deficiencies occuring in plant roots. One of these common responses manifested across multiple nutrient deficiencies in roots was the downregulation of oxidative stress proteins, which play a key signaling role in root development and responses to stress^131^. Previously, ROS accumulation has been observed in plant roots under N, P, and K deficient conditions^132–139^. Specifically, Shin et al. (2005)^136^ identified ROS accumulation occurring primarily in epidermal cells under -N conditions, while under P deficiency, patchier ROS accumulation localized mainly to cortical cells. We observe that N and K deficient conditions had more overlap in ROS scavenging enzymes than P and S deficient conditions, suggesting differential regulation between the nutrient starvation responses at the protein level, which may manifest through different enzymes located in different cell types.

Further, plant root system architecture can be altered under exposure to nutrient deficiency, which also intersects with ROS. Within our root data, we see a downregulation of multiple peroxidases and ascorbate peroxidase. Peroxidases play a key role in modifying cell wall properties^140^. Alteration in peroxidase expression results in altered root growth^141,142^, with peroxidases having been shown to modulate ROS balance at the transition zone between root cell differentiation and proliferation^143^. Our observed downregulation of peroxidases and proteins involved in ROS homeostasis agrees with previous findings examining proteomic changes in canola shoots and roots under N deficiency^73^. Additionally, we find strong downregulation of CATALASE 2 (CAT2) in roots across several nutrient deficiency conditions. CAT2 regulates root meristem activity under oxidative stress^144^, thus its downregulation is likely contributing to nutrient deficiency-dependent changes in root system architecture. Our findings indicate that the dynamics of root ROS scavenging enzymes is broadly affected across multiple nutrient deficiencies, not just -N, with downregulation of protein abundances likely functioning to reshape root system architecture in response to these nutrient deficiencies.

### 4.5 Nutrient deficiency upregulates proteases and protease inhibitors in canola roots

Across all nutrient stress conditions applied here, we uniquely find a general upregulation of proteins involved in peptidase activity and its negative regulation in roots. In particular, papain-like cysteine proteases (PLCPs), RESPONSIVE TO DESSICATION (RD) 21B (duplicates: BnaA06T0471400; BnaC02T0364000) and RD21C (duplicates: BnaC01T0329400; BnaA01T0236000) as well as a protease inhibitor SERPIN1 (duplicates: BnaA10T0058300; BnaC06T0015800), which has been shown to interact with RD21 to inhibit its activity in Arabidopsis seed development, skotomorphogenesis and drought response^145–147^. Thus, our data suggests a novel role for cysteine proteases and inhibitors in canola root responses to nutrient deficiency. We also observe upregulation of another papain inhibitor, CYS1 (duplicates: BnaC09T0508000; BnaA02T0044300; BnaA10T0225700) and a trypsin protease inhibitor KTI1 (BnaC06T0307000), both of which have roles in regulating programmed cell death under oxidative stress and pathogen attack, respectively^148,149^. It is thus possible that these protease inhibitors also serve to regulate and prevent uncontrolled cell death by their upregulation under stressful conditions, which may also be contributing to root system architectural changes.

### 4.6 Calcium signaling proteins play a key role in canola root response to nutrient deficiency

Calcium ions (Ca^2+^) are an important second messenger in response to myriad external stimuli and stresses ranging from growth and developmental cues, to pathogen responses and abiotic stress responses such as temperature, salt and drought^150,151^. The calcium signature hypothesis postulates that different stimuli induce distinct calcium signatures^152–156^, which vary in duration, amplitude, frequency and spatial distribution, and that the unique calcium signature activates appropriate effectors for downstream signaling responses. Calcium signaling has been previously implicated in nutrient deficiency responses, especially in N signaling^157,158^, however there is relatively little known about calcium signaling in P, K and S deficiency signaling. Interestingly, we find an enrichment of ‘calcium binding proteins’ throughout both our GO analysis and STRING-DB association network analysis in roots, noting that many were upregulated across multiple nutrient deficiencies. Further, our data reveals several upregulated calcium-related proteins with a previously uncharacterized function in nutrient deficiency responses. Given the potential pan-nutrient deficiency role that calcium signaling may play, we next chose two calcium signaling proteins for validation. These candidates included: i) CML42, based on its differential regulation in root vs. shoot (up in root, down in shoot), and ii) IQD32 due to its upregulation across all nutrient deficient conditions. Our root growth assays revealed a previously unknown function for CML42 in root response to N deficiency, and for IQD32 in response to S deficiency. It was intriguing that although CML42 and IQD32 showed upregulation across all nutrient deficiency treatments in our proteomic data, they only showed a phenotype under one nutrient starvation stress, suggesting intriguing possibilities relating to more complex regulatory roles and protein networks. For example, CMLs typically act in pairs with their most closely-related homolog^159^, and CML42 and CML43 were both found upregulated across all nutrient stresses in roots our proteomic results. In the case of IQD32, we also identified the closely-related IQD28^160^, similarly upregulated under all nutrient deficiency treatments.

Further analysis of these closely-related proteins would be interesting, but is beyond the scope of the current study. Beyond these two candidates, we also observe differential abundance of CALCIUM-DEPENDENT PROTEIN KINASE 21 (CPK21; BnaA02T0252900), which has been implicated in nitrate signaling to regulate the nitrate transporter, SLAH2/3^161,162^. We also observe differential abundance of a number of additional CPKs and calmodulin-like proteins (CMLs) across multiple nutrient stress conditions, none of which had previously been implicated in nutrient stress signaling. Collectively however, our dataset provides substantial new insights into the seemingly broader role of calcium signaling under nutrient deficient conditions, and sheds light onto a number of putative targets for further analysis that may have higher order nutrient-stress mitigating opportunities for agricultural applications.

## ACKNOWLEDGEMENTS

The authors thank the Natural Sciences and Engineering Research Council of Canada (NSERC) and Canada Foundation for Innovation (CFI) for funding. The authors also thank Jack Moore of the Alberta Proteomics and Mass Spectrometry Facility for assistance with mass spectrometer operation and maintenance.

## AUTHOR CONTRIBUTIONS

LEG: Investigation; Formal Analysis; Methodology; Data curation & Visualisation, Writing & Editing. SS: Investigation; Methodology; Data Curation & Visualization; Writing & Editing.

DM: Investigation; Methodology; Formal Analysis, Data curation & Visualisation IK: Methodology.

RGU: Conceptualization; Methodology; Supervision; Project Administration; Data curation; Writing & Editing; Funding Acquisition.

## DATA AVAILABILITY

The datasets that we have presented in this study are available in public repositories. All proteomic data has been submitted to the PRoteomics IDEntifications Database (PRIDE; https://www.ebi.ac.uk/pride/) with the identification number PXD032223.

## SUPPLEMENTAL DATA

Table S1. Hoagland’s nutrient solution recipes for nutrient deficiency conditions

Table S2. List of all quantified proteins in shoots and roots

Table S3. List of enriched GO terms correlated between two nutrient deficiencies

Table S4. List of duplicated significantly changing proteins represented in association network

Figure S1. Enriched molecular function GO terms Dotplot representation of enriched molecular function GO terms for significantly changing proteins (q-value <0.05; Log2FC >0.58 or <-0.58) that are up- or downregulated in shoot (a) or root (b) under each nutrient deficiency (-N, -P, -K or -S). The size of the dots represents the gene ratio (number of proteins in the conditions/total quantified proteins in the study). The colour code of the dots represents the log10(p-value). Only GO categories with p-value <0.01 and total count <500 are included.

Figure S2. Root growth validation of four candidate proteins across grown on media lacking nitrogen, phosphorus and sulphur Root lengths were measured for wild-type (Col-0) and *cml42*, *hip1, chmp1b* and *iqd32* loss-of-function seedlings 10 days following germination on nitrogen-, phosphorus, and sulphur-deficient media. A two- way ANOVA, followed by Fishers LSD test was used to determine statistical significance between nutrient-deficient conditions and genotypic background. Letters denote statistical significance (<0.05). *cml42*, calmodulin-like protein 42; *iqd32*, IQ-domain 32.

