## Supplemental Figure 1 for "Quantitative proteomic analysis of soil-grown *Brassica napus* responses to nutrient deficiency"

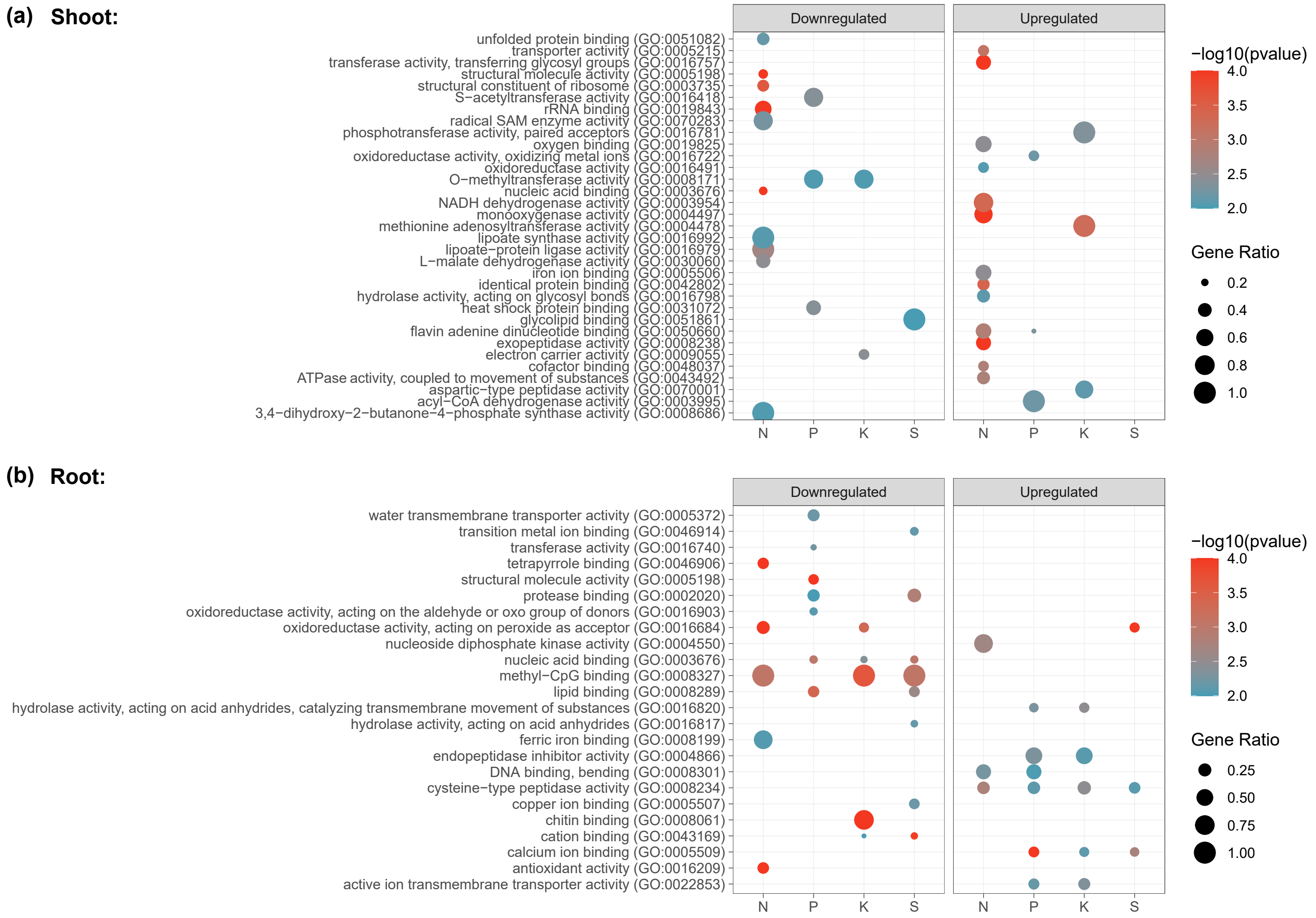


**Figure S1. Enriched molecular function GO terms**

Dotplot representation of enriched molecular function GO terms for significantly changing proteins (q-value <0.05; Log2FC >0.58 or <-0.58) that are up- or downregulated in shoot (A) or root (B) under each nutrient deficiency (-N, -P, -K or -S). The size of the dots represents the gene ratio (number of proteins in the conditions/total quantified proteins in the study). The colour code of the dots represents the log10(p-value). Only GO categories with p-value <0.01 and total count <500 are included.
