## Supplemental Figure 2 for "Quantitative proteomic analysis of soil-grown *Brassica napus* responses to nutrient deficiency"

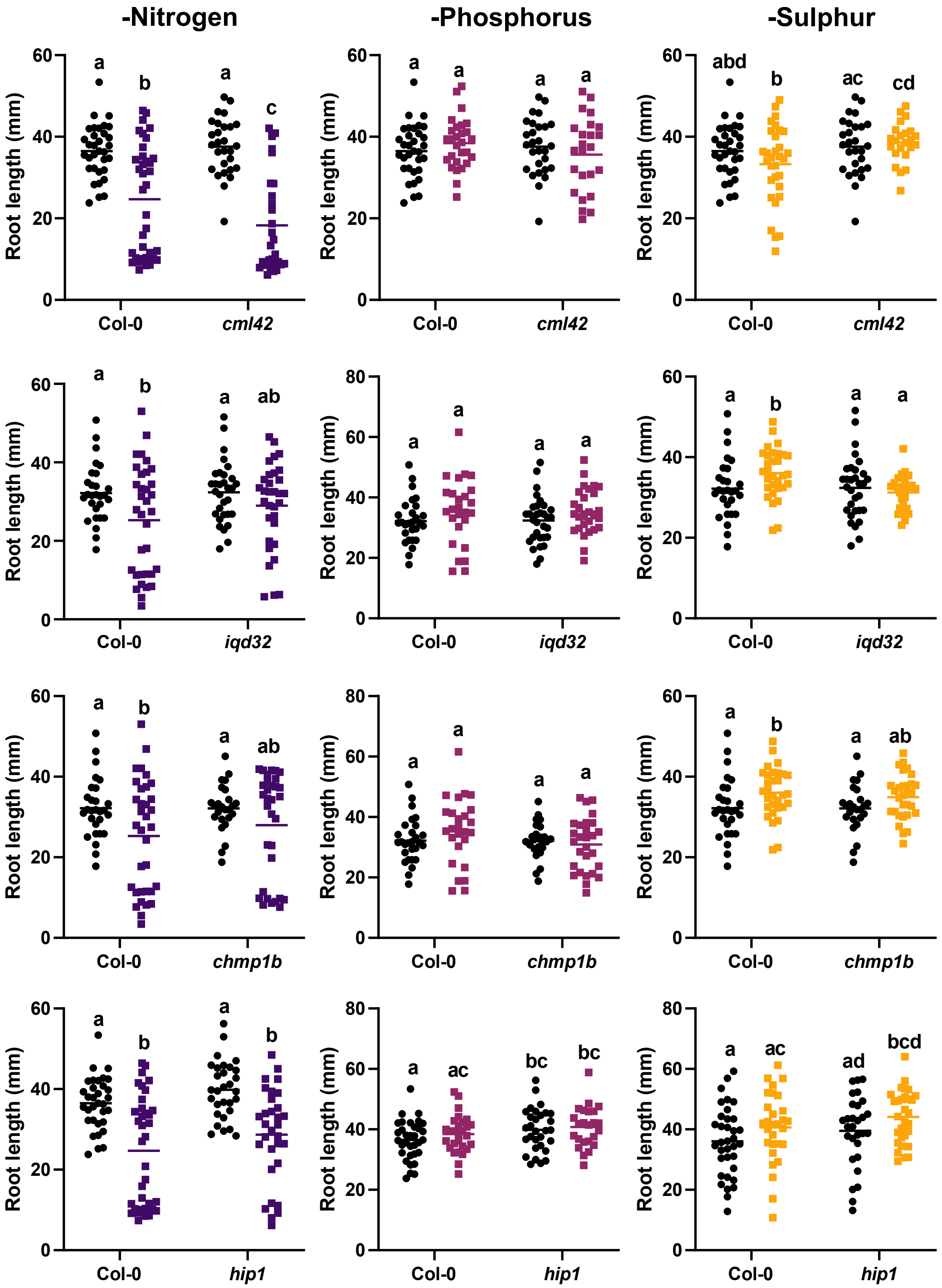


**Figure S2. Root growth validation of four candidate proteins across grown on media lacking nitrogen, phosphorus and sulphur**

Root lengths were measured for wild-type (Col-0) and *cml42*, *hip1, chmp1b* and *iqd32* loss-of-function seedlings 10 days following germination on nitrogen-, phosphorus, and sulphur-deficient media. A two-way ANOVA, followed by Fishers LSD test was used to determine statistical significance between nutrient-deficient conditions and genotypic background. Letters denote statistical significance (<0.05). *cml42*, calmodulin-like protein 42; *iqd32*, IQ-domain 32.
